# *Tribolium castaneum* as a model for microRNA evolution, expression and function during short germband development

**DOI:** 10.1101/018424

**Authors:** Maria Ninova, Matthew Ronshaugen, Sam Griffiths-Jones

**Affiliations:** Faculty of Life Sciences, Michael Smith Building, Oxford Road, University of Manchester, Manchester, M13 9PT, UK

**Keywords:** microRNA clusters, short germband development, maternal-zygotic transition, maternal transcripts

## Abstract

MicroRNAs are well-established players in the development of multicellular animals. Most of our understanding of microRNA function in arthropod development comes from studies in *Drosophila.* Despite their advantages as model systems, the long germband embryogenesis of fruit flies is an evolutionary derived state restricted to several holometabolous insect lineages. MicroRNA evolution and expression across development in animals exhibiting the ancestral and more widespread short germband mode of embryogenesis has not been characterized. We sequenced small RNA libraries of oocytes and successive time intervals covering the embryonic development of the short germband model organism, the red flour beetle *Tribolium castaneum.* We analysed the evolution and temporal expression of the microRNA complement, and sequenced libraries of total RNA to investigate the relationships with microRNA target expression. We show microRNA maternal loading and sequence-specific 3’-end non-template oligoadenylation of maternally deposited microRNAs that is conserved between *Tribolium* and *Drosophila.* We further uncover large clusters encoding multiple paralogs from several *Tribolium-specific* microRNA families expressed during a narrow interval of time immediately after the activation of zygotic transcription. These novel microRNAs, together with several early expressed conserved microRNAs, target a significant number of maternally deposited transcripts. Comparison with *Drosophila* shows that microRNA-mediated maternal transcript targeting is a conserved phenomenon in insects, but the number and sequences of microRNAs involved in this process have diverged. The expression of fast-evolving and species-specific microRNAs in the early blastoderm of *T. castaneum* is consistent with previous findings in *Drosophila* and shows that the unique permissiveness for microRNA innovation at this stage is a conserved phenomenon.

## Introduction

MicroRNAs are short non-protein coding RNAs, processed from hairpin precursors. MicroRNAs regulate gene expression by guiding the RNA-induced silencing complex (RISC) to complementary sites in the 3’ UTRs of target mRNAs, thereby inducing translational silencing and degradation (reviewed in Bartel 2004). MicroRNA-target interaction usually requires base pairing within a 6-7-mer “seed” region starting from the second nucleotide from the 5’ end of the mature product, and thus each microRNA can potentially target hundreds of protein-coding genes (reviewed in Bartel 2009). Consistent with their functional importance, seed sequences are the most highly conserved regions of the microRNA hairpins (Lim et al. 2003b, 2003a; Lai et al. 2003). MicroRNAs were first identified for their role in the regulation of developmental timing in *Caenorhabditis elegans* (Lee et al. 1993; Reinhart et al. 2000), and were later shown to play important roles in various aspects of the development of both invertebrates and vertebrates (reviewd in Kloosterman and Plasterk 2006). Most of our understanding of microRNA function in the development of arthropods comes from studies in the classical model organism *Drosophila melanogaster.* MicroRNAs in *D. melanogaster* have been shown to control essential developmental processes such as the maternally deposited transcript clearance, cell differentiation and apoptosis, morphogenesis and organogenesis (Asgari 2013).

In general, arthropods share a common conserved anatomical architecture – the segmented body – that is established during the early stages of embryogenesis. However, there is significant variation between animals in the developmental mechanisms that precede and follow this segmented body stage. *Drosophila* development follows the so-called ‘long germband’ developmental mode, which is derived and found only in a subset of holometabolous insect lineages (Peel 2008; Mito et al. 2010). The most common, and likely ancestral, mode of arthropod pre-segmentation developmental is short germband embryogenesis (Peel 2008; Mito et al. 2010) In short germband embryogenesis, a small number of cells in the blastoderm embryo form the most anterior segments (called the germ anlage), while more posterior segments arise after gastrulation by growth and cell division (reviewed in Davis and Patel 2002). Whereas long germ development often occurs in a syncytium, short germ segmental patterning occurs in a cellularized environment via an oscillatory mechanism (Sarrazin et al. 2012). The remaining portion of the early blastoderm gives rise to the extra-embryonic serosal membrane (Handel et al. 2000). In many respects, short germband embryogenesis therefore more closely resembles the segmentation of vertebrate embryos.

The red flour beetle *Tribolium castaneum* is an emerging model organism that displays a number of ancestral features, including the short germband mode of development (Richards et al. 2008; Roth and Hartenstein 2008; Denell 2004). The availability of genetic tools and a wide range of embryonic patterning mutants have established *T. castaneum* as a model system to study this ancestral developmental mode (Richards et al. 2008; Denell 2004). *T. castaneum* has a fully sequenced genome (Richards et al. 2008) and an annotated protein-coding transcriptome (Kim et al. 2010). In addition, the morphology of its early embryogenesis is among the best characterized of the short germband insects (Handel et al. 2005, 2000; Denell 2004; Benton et al. 2013).

MicroRNAs are recognised as important players in developmental gene regulation, yet their expression and function in short-germband embryogenesis is poorly understood. Furthermore, little is known about the evolutionary constraints that act on microRNA developmental expression in general. We explored the microRNA complement of *T. castaneum* and expression of their targets in a developmental context, using a combination of small RNA and whole transcriptome RNA sequencing of successive time intervals covering beetle embryogenesis. We report significant amounts of maternally-deposited microRNAs in *T. castaneum* oocytes, and conserved non-template oligoadenylation of specific mature microRNA sequences. During embryogenesis, microRNA abundance markedly increases at the onset of zygotic transcription, including the vast majority of conserved microRNAs, as well as a large number of previously-unannotated microRNAs organized in multiple rapidly-evolving multicopy clusters. Expression of these rapidly-evolving microRNAs is uniquely upregulated during a very narrow period during blastoderm cleavage. We show that maternally-deposited protein-coding mRNAs that are down-regulated in the early embryo are significantly enriched in targets of novel and conserved microRNA families that are up-regulated in this narrow developmental window. We therefore show a role for early expressed microRNAs in maternal transcript clearance in *Tribolium.* Comparisons with previous findings of microRNA-mediated maternal transcript degradation in *Drosophila* (Bushati et al. 2008) reveal conserved and divergent aspects of this process in holometabolous insects. Taken together, these results provide an unprecedented insight into the evolution, expression and function of microRNAs in the early stages of arthropod development.

## Results

### *T. castaneum* small RNA sequencing and annotation

*T. castaneum* embryonic development is significantly longer than that of fruit flies, spanning 6 days at 25°C. We collected embryonic samples from different time intervals after egg laying to determine the time points when key developmental events occurred (Supplemental Text; Supplemental Figure 1). Based on these observations, we generated small RNA sequencing libraries from seven discrete intervals of *T. castaneum* embryogenesis covering the following key stages: very early embryo before the onset of zygotic transcription (0-5 h, herein referred to as “pre-ZT” embryos; note that zygotic transcription occurs at ∼8 h, Supplemental Figure 1B); later cleavage divisions and blastoderm formation (8-16 h); blastoderm differentiation and beginning of gastrulation (16-20 h); progressing serosal closure (20-24 h); elongating germband (24-34 h); fully segmented germband and appendage formation onset (34-48 h); extended germband until hatching (48−144 h). Accurate staging was verified by microscopic inspection of an aliquot of each sample (see below). We obtained between 3.5 and 6.5 million reads for each sample, over 85% of which mapped to the 10 chromosomes or the unassembled scaffolds of *T. castaneum* genome with no more than 1 mismatch.

The previously-annotated set of microRNAs in *T. castaneum* comprises 203 hairpins that were experimentally identified in mixed adult and mixed embryonic datasets. Nearly half of these sequences have no identifiable orthologs in other species with sequenced genomes (Marco et al. 2010, Supplemental Table 1). We took advantage of the discrete embryonic stage datasets to search for putative novel microRNAs that may be expressed during narrow periods of time, and thus escaped previous detection. Using two independent approaches for novel microRNA identification from deep sequencing data, we revised previous annotations and uncovered a total of 124 novel *Tribolium-specific* microRNA candidates (see Supplemental Table 1). A large fraction of these hairpins are paralogs of, or have similar extended seed sequences to the microRNA families mir-3851 and mir-3836. Homology searches suggest that mir-3851 family is *Tribolium-specific.* The miR-3836-3p sequence has 13 nt identity with the *Drosophila* miR-3-3p sequence, and an identical 8mer region including the seed sequence with miR-309-3p. However, there is no other obvious sequence similarity between the hairpins, and thus homology cannot be confidently assigned. Regardless of their ancestral origins, the large number of paralogs from the mir-3836 family in *T. castaneum* suggests that they emerged by multiple lineage-specific duplications. Furthermore, there is a second set of novel sequences that lack homology with any known microRNAs, but are found in multiple copies and can be grouped in several families. Sequence alignments of the mir-3851, mir-3836 and novel families are shown in Supplemental Figure 2. Notably, the vast majority of mir-3851, mir-3836 and novel microRNA family members are organized in clusters in the genome, mostly localized in subtelomeric regions or unmapped scaffolds (Figure 1). Even though these clusters encode homologous microRNAs from several families, they differ in their members’ copy number, sequence and organization, suggesting rapid evolution by duplication and diversification. We collectively refer to these clusters as “multicopy microRNA clusters”.

**Figure 1.**
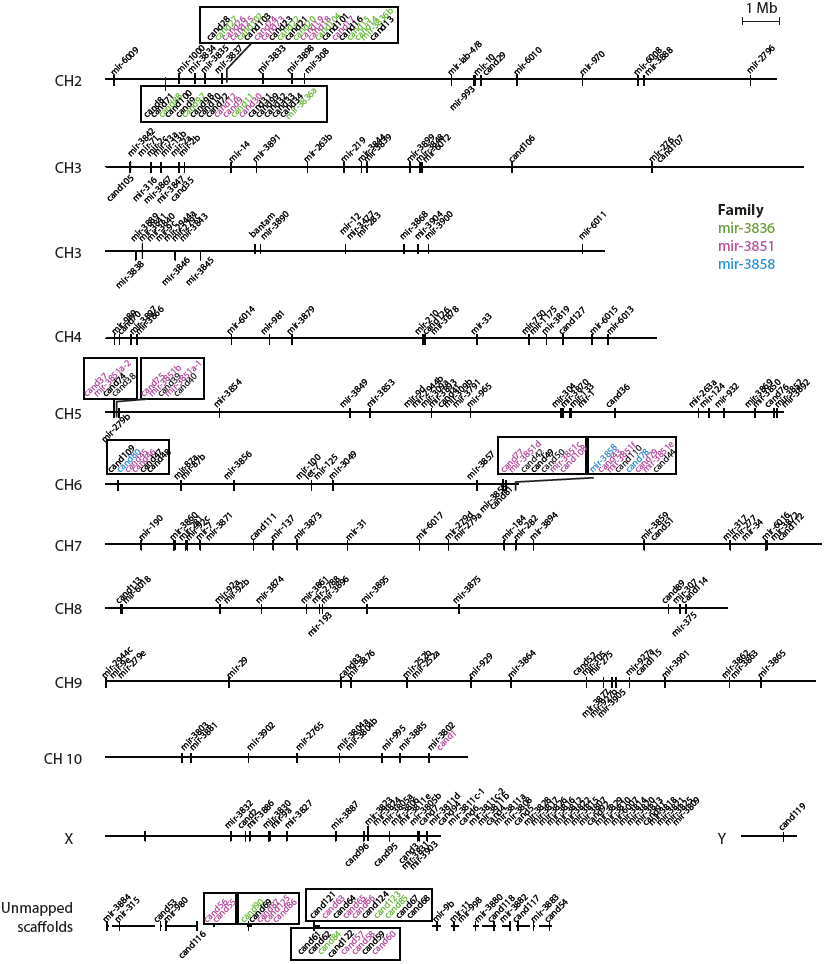
Genomic organisation of known and novel candidate microRNAs in *T. castaneum*. Horizontal lines represent the assembled chromosomes and the unmapped scaffolds of the *T.castaneum* genome assembly (r4.0), on which microRNA genes are found. Different microRNA-encoding loci are marked, and microRNAs within a 10 Kb distance from each other are shown together as a cluster. ‘Cand’-number denotes candidate novel microRNAs. Different colours highlight members of three widespread *Tribolium-specific* families of microRNAs, and black boxes highlight clusters encoding mir-3836 and mir-3851 family members.

### Small RNA temporal dynamics during *T. castaneum* embryogenesis

To gain insight into the developmental dynamics of small RNAs during *T. castaneum* development, we first assessed the overall content of the small RNA sequencing libraries of successive time intervals. Figure 2 shows the distributions of the absolute counts of small RNA reads of sizes from 17 to 35 nt in the corresponding embryonic stage; the number of reads of each size that map to up to 5 positions in the genome annotated as microRNAs, protein-coding genes, tRNAs, or intergenic regions; reads that map to multiple (>5) positions; and those that cannot be mapped. Reads from all libraries and sizes display a very strong bias for uracil in the first position, which is typical for microRNAs and piRNAs. Size distributions of the sequenced reads display two peaks at ∼22 nt and ∼28 nt, with the vast majority of the ∼22 nt reads corresponding to known or newly annotated microRNAs. The ∼28 nt fraction at all stages comprises highly heterogeneous reads originating from diverse genic and intergenic regions, both unique and repetitive loci. Manual inspection of the read positions and patterns strongly suggests that these reads mostly represent primary and secondary piRNAs that are maternally deposited or zygotically transcribed in response to active transposable elements in the embryo (manuscript in preparation). The relative abundances of putative microRNAs and piRNAs change markedly as development progresses. piRNAs are highly abundant in the earliest stages of development, consistent with the notion of maternal deposition of this class of molecules (Brennecke et al. 2008). The relative microRNA levels on the other hand are initially very low; the ∼22 nt peak is almost absent prior to activation of zygotic transcription. The microRNA/piRNA ratio then increases with developmental time, until microRNAs account for nearly 30% of all small RNA reads in the late embryo. These figures are in sharp contrast with the small RNA distributions observed in early embryonic developmental datasets from *Drosophila* previously generated by our group and others, where microRNAs represent the most significant peak in early embryonic small RNAs (see below), and dominate the non-ribosomal small RNA fraction past the first couple of hours of development.

**Figure 2.**
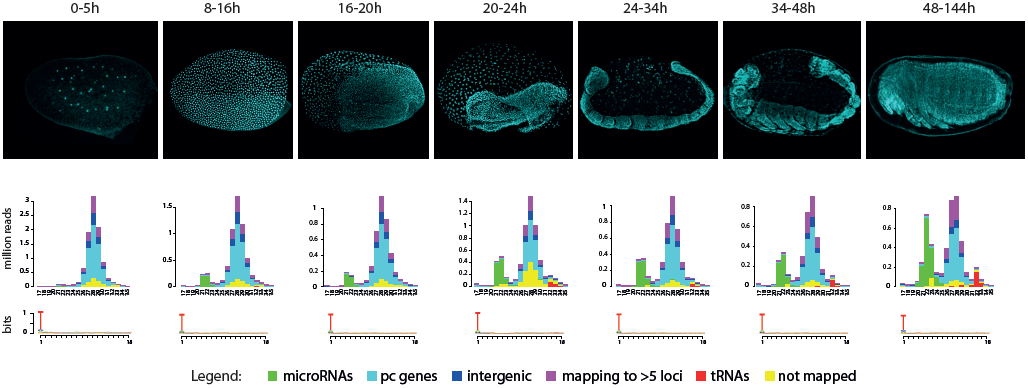
Small RNA types and size distributions throughout *T. castaneum* development. Histograms show the small RNA size and count distribution in each small RNA library (DAPI-stained representative images for the corresponding time intervals are shown in the top panel). Different colours reflect read mapping status: not mapped, mapping to >5 loci or mapping to at least one tRNA, microRNA, protein coding (pc) gene or intergenic regions. Sequence logos show the nucleotide bias for the first 18 positions of reads.

### Abundance, diversity, conservation and 3’-end modification of maternally deposited microRNAs

We hypothesized that the apparent low microRNA levels in *T. castaneum* pre-ZT embryos reflects either insignificant maternal loading of microRNAs in oocytes, or decreased sampling of microRNAs by sequencing due to the highly abundant piRNA fraction. We therefore sought to determine the absolute amount of microRNAs deposited in *Tribolium* and *Drosophila* eggs. To this end, we sequenced the small RNAs of samples prepared from a fixed number of unfertilized eggs from *T. castaneum* and two divergent fruit fly species, *D. melanogaster* and *D. virilis*, with spiked-in synthetic 5’-phosphorylated oligonucleotides at different final concentrations ranging between 0.1 and 0.001 fmol per cell (see Materials and Methods). In addition, we used qPCR to quantify the absolute cellular levels of miR-184-3p, an abundant microRNA that is perfectly conserved between the three species, and used this quantity as an additional endogenous reference. miR-184-3p quantities determined independently by a standard curve and relative to the 0.1 fmol spike-in (cel-mir-35-3p) were similar; further, the miR-184-3p:cel-mir-35-3p ratios in deep sequencing read counts were in a good agreement with the qPCR estimates, confirming that the sequencing data reflect well the microRNA abundances in this concentration range (Supplemental Text and Supplemental Figure 3). The lower concentration spike-ins had very few reads in the deep sequencing experiments and were therefore ignored for further analysis. Small RNA size distributions of the resulting libraries are shown in Figure 3A. Similarly to the transcriptionally inactive 0-5 h embryos, *T. castaneum* oocytes are characterized by abundant 28 nt sequences, consistent with piRNAs, and relatively low microRNA levels. In contrast, *D. melanogaster* and *D. virilis* oocyte small RNA profiles display two prominent peaks corresponding to maternally provided microRNAs and putative piRNAs. Normalizing the microRNA read counts according to the known concentrations of miR-184-3p and the spiked-in reference independently and consistently gave an estimate of approximately 0.2 fmoles of microRNAs per egg in *Tribolium*, and approximately 4 times higher microRNA content in *Drosophila* (Figure 3B,C). This equates to an estimate of ∼120 million individual microRNA molecules per *Tribolium* egg. Previous studies suggest that these levels are likely to be physiologically meaningful. For example, previous microRNA estimates in mammalian hematopoetic cells suggested that the most highly expressed sequences are found at ∼1000-2000 copies per cell, with a median of 178 (Bissels et al. 2009). However, we note that a 4-fold difference in microRNA abundance between the fly and beetle is not sufficient to explain the difference in the microRNA/piRNA ratios in the two taxa.

**Figure 3.**
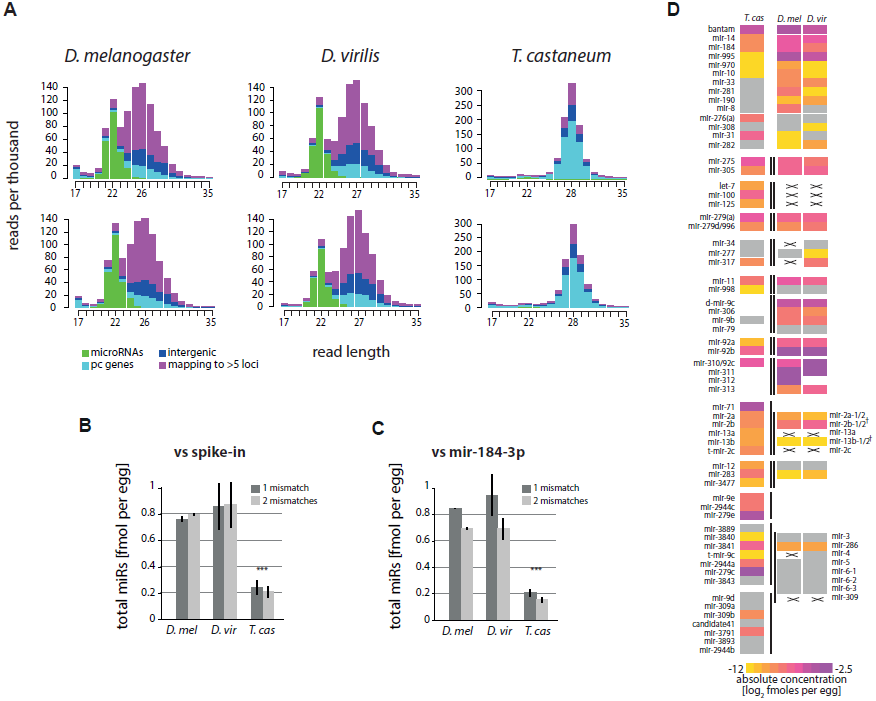
Quantities and conservation of maternally deposited microRNAs. A. Read size and count distributions of small RNA sequencing libraries from oocytes in the three species (two biological replicates per species). Reads were coloured depending on their mapping position to microRNAs, protein-coding (pc), intergenic regions or multiple (>5) sites in the genome. B. Absolute abundance of total microRNAs as determined by deep sequencing read counts relative to spike-in. C. Absolute abundance of total microRNAs as determined by deep sequencing read counts relative to miR-184-3p. B, C. Quantification from deep sequencing analysis was performed using 1 and 2 allowed mismatches between read and genome. D. Heat map showing the estimated concentrations of conserved microRNAs or clusters of microRNAs between Drosophilids and *T. castaneum.* MicroRNAs below an arbitrary threshold of 0.0002 fmol/cell are shown in grey. Lines mark clustered microRNAs. Crosses mark conserved microRNAs that are not expressed, while blank spaces denote absence of a given homolog from the corresponding cluster in a given species. MicroRNAs are aligned by homology; note that mir-3∼309 cluster of *Drosophila* has three homologs in *Tribolium.* ^†^mir-2 cluster in *Drosophila* has split.

We next assessed the most abundant maternally deposited microRNAs in *Tribolium*, and compared these to fruit flies. Figure 3D shows the expression of conserved singleton microRNAs and clusters of microRNAs with at least one conserved member between the two taxa (for full read counts see Supplemental Table 2). The data show that the conserved microRNA families bantam, mir-275, mir-305, mir-14, mir-184, mir-995, mir-2/11/13, mir-92/310, mir-279 and mir-9 represent the most highly maternally loaded microRNAs in oocytes of both fruit flies and the flour beetle. For clustered microRNAs, either all or none of the members of a cluster are usually present in the oocyte, consistent with the notion that clustered microRNAs are co-expressed (Baskerville and Bartel 2005; Ruby et al. 2007; Ryazansky et al. 2011). A notable example of divergence in terms of maternal deposition is the mir-100/let-7/mir-125 cluster, which are loaded in *Tribolium* oocytes, but not in *Drosophila.* In addition, the most abundant microRNAs in the beetle oocyte belong to two clusters; *mir-279e∼2944c* and *mir-3889∼3843.* These are diverged homologs of the *Drosophila mir-309∼6* and *mir-994∼318* (Ninova et al. 2014), whose maternal deposition is *Drosophila* is modest. The maternal microRNA complement of *T. castaneum* also contains a number of lineage-specific microRNAs, including the members of the massively duplicated 40-mir cluster on the X chromosome, albeit at lower concentrations (Supplemental Table 2).

MicroRNA 3’-ends are known to be subject to various modifications in different model systems, including ligation of additional nucleotides to the 3’-end of the mature product (Ha and Kim 2014). In *D. melanogaster*, up to 5% of the microRNAs were previously found to contain non-templated adenosine or uracil residues at their 3’-end in early embryonic samples (Fernandez-Valverde et al. 2010), and a recent study reported even higher levels of 3’-end modifications in oocytes and early embryos of *D. melanogaster* and deuterostomes (Lee et al. 2014). 3’-end non-template modification of oocyte microRNAs in *D. melanogaster* is mediated by the non-canonical poly-A polymerase Wispy, and this modification was suggested to be involved in maternal microRNA clearance (Lee et al. 2014), but the precise role of microRNA tailing in fly development is not well understood. To address the presence and developmental dynamics of microRNA 3’-end nucleotide additions in *T. castaneum*, we developed a computational pipeline that searches for reads that perfectly map to the dominant isoforms of the microRNA mature arms, but have additional nucleotides that do not match the genome at the 3’-end. In addition, we analysed *D. virilis* developmental time series previously generated by our group for comparison (Ninova et al. 2014). The data show that 3’-end non-template additions affect at least 20% of the maternally deposited microRNAs (Figure 4A), which is significantly higher than later times or in other tissues, and is consistent with the previous observations in *D. melanogaster* (Lee et al. 2014a). The majority of non-template 3’-end additions are between one and three nucleotides in length (Figure 4B), and are almost exclusively adenosines (Figure 4C). We note that some microRNAs have an adenosine in the genomic position immediately following the predicted 3’-end Drosha cleavage site, thus post-processing adenylation is impossible to distinguish from imperfect processing, and we therefore exclude such cases from the analysis. Our estimates of the extent of microRNA modifications in insect oocytes are therefore likely to be conservative. The proportion of reads with modified ends varies between different microRNAs and is poorly correlated with expression level – while some highly abundant microRNAs are not significantly modified, several highly abundant microRNAs display a particularly high ratio between their modified and unmodified forms in oocytes, including some microRNAs that are specifically expressed in a given species, such as the miR-100/let-7/miR-125 in *T. castaneum.* (Figure 4D). This observation is consistent with recent data in *D. melanogaster* by Lee and colleagues, and the same selected microRNA species appear to be strongly modified in all *D. melanogaster* libraries (Lee at al. 2014). Amongst the conserved microRNAs maternally deposited at high levels in both *Drosophila* species and *Tribolium* products of the same families (e.g. miR-184-3p, miR-279-3p, miR-9-3p, miR-92/310-313-3p) are heavily modified. Thus, the process of oligoadenylation of specific maternally deposited microRNAs is conserved between the two insect taxa.

**Figure 4.**
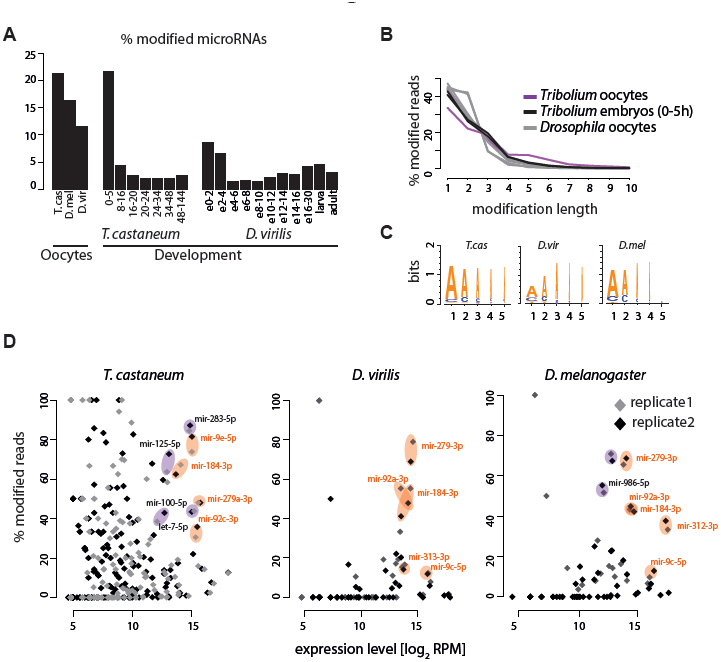
Non-templated 3’-end nucleotides in oocyte microRNAs. A. Proportion of microRNAs with non-template 3’-end nucleotide addition in oocytes and developmental small RNA libraries of *T. castaneum* and *Drosophila*. B. Length distribution of non-template 3’-end nucleotide additions. C. Sequence logos showing the most frequent nucleotides in the first 5 positions of the 3’-end non-templated nucleotide additions in the three species. D. Scatter plots showing the proportion of reads with non-template 3’-end additions as a function of the total read counts (modified and non-modified) per million for each microRNA. Results of different replicates for each species are shown separately. Example singleton microRNAs or microRNAs from clusters that are conservatively deposited in the egg are highlighted in orange, and highly deposited and strongly modified microRNAs specific for a given species are highlighted in purple.

### Developmental expression of conserved and non-conserved microRNAs

We next investigated the temporal dynamics of microRNA expression throughout beetle development. Full read counts of all mature microRNAs from the different small RNA sequencing datasets are shown in Supplemental Table 2. Figure 5A shows the correlations of microRNA expression profiles between the different developmental stages of *Tribolium* in an all-versus-all manner. As expected, the microRNA repertoire in oocytes and 0-5 h embryos is highly similar, as zygotic expression is not active during the initial cleavage divisions. The similarity between pre-ZT and blastoderm embryos (8-16 h) however is significantly lower, indicating a shift in the microRNA expression profile upon activation of zygotic transcription. Subsequently, the correlation of microRNA expression is the highest between neighbouring stages, and similarity decreases with increasing developmental distance. Thus, developmental transitions in *T. castaneum* are accompanied by shifts in the global microRNA profiles.

**Figure 5.**
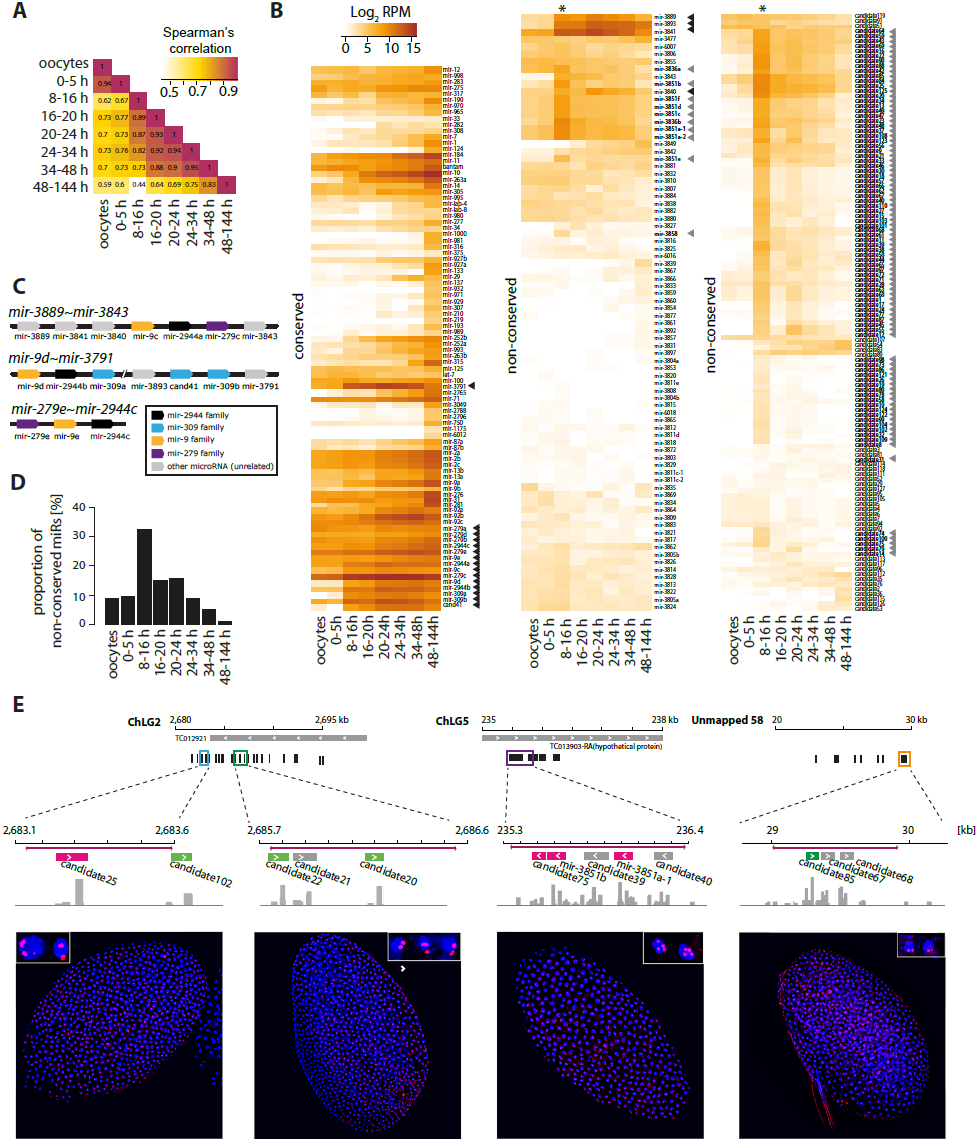
Developmental expression of conserved and non-conserved microRNAs throughout *T. castaneum* development. A. Heatmaps representing Spearman’s correlation values for all-versus-all comparisons of the microRNA expression levels in oocytes and embryonic intervals of 0-5 h, 8-16 h, 16-20 h, 20-24 h, 24-34 h, 34-48 h, and 48-144 h. B. Heatmaps showing the normalized expression levels (reads per million) of conserved and species-specific microRNAs in *T. castaneum* oocytes and developmental intervals. Grey arrowheads indicate microRNAs from the mir-3851 and mir-3836 families and other microRNAs clustered in the same loci as in Figure 1. Black arrowheads mark members of the *mir-279e∼2944c*, *mir-3889∼3843* and *mir-9d∼3791* clusters. C. Diagram of the genomic organization of the *T. castaneum mir-279e∼2944c*, *mir-3889∼mir-3843* and *mir-9d∼3791* clusters. MicroRNAs are color-coded based on sequence homology. D. Relative proportion of *Tribolium-specific* microRNA reads at different developmental stages. E. Spatial expression of *Tribolium-specific* microRNA clusters detected by *in situ* hybridization. Top diagrams indicate the genomic regions encoding *Tribolium-specific* microRNAs used for antisense DIG-labelled RNA probe design, with probe regions in coloured boxes. MicroRNA genes are black or colour-coded based on homology to mir-3851 (magenta) and mir-3838 (green). Histograms represent coverage tracks generated with IGV. (Bottom) Confocal images of *T. castaneum* blastoderm embryos showing nascent microRNA transcripts (red); blue represents DAPI nuclear staining.

To gain a further insight in the diversity and dynamics of the microRNA complement throughout beetle embryogenesis, we assessed the normalized expression levels of the annotated *T. castaneum* microRNAs at each developmental interval. Heatmaps in Figure 5B show the levels of individual microRNAs grouped by conservation in other species. Consistent with previous notions that conserved microRNAs are more highly expressed (Roux et al. 2012; Meunier et al. 2013; Liang and Li 2009; Ruby et al. 2007), conserved microRNAs dominate the expression profiles in all embryonic stages of *T. castaneum.* We detect expression of virtually all annotated conserved microRNAs during at least one stage of development, with the vast majority displaying their highest relative levels in the late embryo during morphogenesis and organogenesis (48 h – 6 days). The most strongly expressed microRNAs during the early and intermediate stages of embryogenesis derive from three clusters, *mir-9d∼3791*, *mir-3889∼3843* and *mir-279e∼2944c* (Figure 5B, black arrowheads). These clusters encode a combination of conserved and *Tribolium*-specific microRNAs (Figure 5C); clusters encoding homologs of the conserved mir-9, mir-2944, mir-279, mir-3791 and mir-309 families are found in other insect lineages, and represent one of the most extreme examples of microRNA gain, loss, duplication, and rearrangement reported to date (Ninova et al. 2014). Previous studies by us and others have shown that members of one of these clusters (*mir-309∼6*) in *Drosophila* are strongly upregulated in the early fly blastoderm (Biemar et al. 2005; Ninova et al. 2014), and that members of the homologous clusters in *A. mellifera* (*mir-3478∼318*) and mosquitoes (*mir-309∼286*) are also highly expressed in early embryos (Zondag et al. 2012; Hu et al. 2014). In addition, the mosquito-specific *mir-2941∼2946* cluster, whose 3’ mature sequences have the same seed as miR-3889-3p in *T. castaneum*, are the most abundant microRNAs in the early stages of development (Hu et al. 2014). We detect nascent primary transcripts of *mir-9d∼3791* and *mir-3889∼3843* in the early blastoderm nuclei of *T. castaneum* by *in situ* hybridization (Supplemental Figure 4A). Taken together, these data suggest that, despite multiple rearrangements, the early zygotic onset of expression of these groups of microRNAs is conserved. Interestingly, we detect nascent *mir-9d∼3791* transcripts much later in development, in serosal nuclei, suggesting a novel role of members of this cluster (Supplemental Figure 4B).

While most stages of development are characterized by low levels of *T. castaneum-specific* microRNAs, the majority of non-conserved microRNAs display a coordinated sharp increase in expression at the undifferentiated blastoderm stage (8-16 h), suggesting that the onset of zygotic transcription is accompanied by up-regulation of multiple non-conserved microRNAs (Figure 5B, asterisk). Indeed, the early blastoderm shows the greatest relative expression level of non-conserved microRNAs across all tested developmental stages (Figure 5D). Assessment of the genomic loci and microRNA families showed that without exception, the blastoderm-specific pool of non-conserved microRNAs belong to the multicopy novel clusters encoding divergent members of the mir-3851, mir-3836 and other *Tribolium-specific* microRNA families described above (Figure 5B, grey arrowheads, also see Figure 1). Their expression patterns suggest that these diverse multicopy microRNA clusters are up-regulated for a discrete period of time, immediately following the onset of zygotic transcription in the blastoderm and are rapidly extinguished shortly afterwards. We generated fluorescently labelled antisense RNA probes against ∼1 Kb regions overlapping the most highly expressed microRNAs from different regions and detected nascent transcripts by *in situ* hybridization (Figure 5E). Due to the high sequence similarity between some regions, we expect that a subset of probes would cross-hybridize and detect transcription from more than one locus, and indeed this is the case for the mir-3851a-1 region (Figure 5E, third panel). The data show that multicopy microRNA clusters are ubiquitously expressed in all blastoderm nuclei, but surprisingly, their expression is limited to a very narrow time interval from the 8-9^th^ to the 11^th^ cleavage division. We conclude that the multicopy microRNA clusters are largely silenced after this time point.

### Abundant early expressed microRNAs target maternally loaded and zygotically down-regulated genes

Very early expressed microRNAs have been shown to play a role in the clearance of maternally deposited transcripts in both *Drosophila* and zebrafish (Giraldez et al. 2006; Bushati et al. 2008). However, the microRNAs involved in these process in the two species (*mir-309∼6* and *mir-430* clusters, respectively), are not homologous, and microRNA-dependent maternal transcript clearance is thought to be a convergent phenomenon. Nonetheless, a common feature is that in both taxa these microRNAs are encoded in large clusters that have undergone multiple duplications and diversification. The most highly expressed microRNAs in the early embryo of *T. castaneum* are also encoded in large clusters, including *mir-9d∼3791* (homologous to the fruit fly *mir-309∼6* cluster (Ninova et al. 2014)) *mir-279e∼2944c*, and *mir-3889∼3843* (Figure 5B,C), as well as other large species-specific clusters discussed above (Figure 1, Figure 5B). Thus, we asked whether these or other *T. castaneum* microRNAs might be involved in maternal transcript clearance.

To determine the maternally deposited mRNA complement, and its fate after zygotic genome activation, we used RNA sequencing to estimate protein-coding transcript levels in unfertilized oocytes, early blastoderm embryos (8-16 h), embryos at the stage of blastoderm differentiation, gastrulation and serosal closure (16-24 h), and embryos at the stage of germband elongation and segmentation (24-48 h) (data deposited in GEO with accession GSE63770). The resulting ∼300 million paired end reads were mapped against the *T. castaneum* genome and transcriptome, detecting at least one fragment for 15221 out of the 16503 annotated protein-coding genes (92%). Gene expression levels were highly similar between replicates (r > 0.96), and differential expression analyses show that a large number of transcripts significantly change their abundance between different intervals (Figure 6A,B). In particular, we find a large number of transcripts that are highly up-regulated between embryonic development and oocytes, likely reflecting developmental processes activated in the early embryo. A substantial number of transcripts also display a smaller yet significant negative change in their levels with the progression of embryogenesis. We next predicted the putative microRNA targets of the *T. castaneum* transcripts by two different microRNA target prediction approaches – detection of canonical microRNA-target binding sites (Bartel 2009), and using the miRanda algorithm, which takes into account sequence complementarity and RNA-RNA duplex free energy (Enright et al. 2003). These methods resulted in 12565 and 12660 predicted microRNA targets among the 13412 genes with available 3’UTR annotations, respectively, with 144579 individual microRNA-target pairs overlapping between the two sets, and 276412 and 241723 pairs unique for each set. We then calculated whether the targets of individual mature microRNAs are enriched among the protein-coding transcripts that are significantly (more than 2-fold) down-, up- or not regulated between oocytes and embryos. Despite the poor overlap of individual target sites between the target prediction methods, the overall trends in microRNA target enrichment are consistent. Distributions of the hypergeometric p-values of target enrichment based on canonical interactions are shown in Figure 6C, and the complete datasets for both target prediction algorithms, as well as analyses performed using an alternative differential expression algorithm, are available as Supplemental Table 3. The data show that genes that are down-regulated between oocytes and embryos, particularly in the early blastoderm (8-16 h), are strongly enriched in targets of a specific set of microRNAs. In general, these microRNAs do not target a significant proportion of genes that are up-regulated or that maintain their expression. We further assessed the expression levels, sequence, genomic localization and evolutionary relationships of the microRNAs that specifically target zygotically down-regulated genes but not up-regulated genes (enrichment of targets in the down-regulated set with p < 0.001 after Bonferroni’s correction). First, we note that this set of microRNAs consists of multiple paralogs from a small number of families, inferred by manual inspection of hairpin multiple sequence alignments (Figure 6C labels; Supplemental Figure 2). Most representatives of these families are clustered in the genome in various configurations, including both *T. castaneum-specific* clusters and clusters with homologs in other species (see Figure 1). Altogether, this set of microRNAs includes 15 of the 16 members of the *mir-279e∼2944c*, *mir-3889∼3843* and *mir-9d∼3791* clusters, as well as novel and previously annotated members of the mir-3851 and mir-3836 families, which are uniquely up-regulated upon the activation of zygotic transcription. Sequence comparisons further showed that many microRNAs that lack overall sequence similarity have identical 6-,7- or 8-mer seed regions (herein referred as “seed families”; Figure 6D), thus have highly similar predicted canonical target sets. Note that seed and evolutionary related families do not always fully overlap: there are microRNAs without detectable common evolutionary origins but with identical seeds, but also evolutionarily related members of the mir-3851 family seed sequences have diverged by one nucleotide and their predicted target sets differ. Major seed families include AGUACG (3p arms of mir-3851b-f, mir-3889, mir-3840, and mir-3841), AGUACA (3p arm of mir-3851a), CACUGG (3p arms of mir-3836 and mir-309/3 families), AUCACA (3p arms of mir-2944, mir-2/13, mir-11 and mir-308; complementary to the previously described K-box motif, Lai et al. 1998), UAAAGC (3p arms of mir-9, mir-3893 and mir-3843; complementary to the Brd-box motif, Leviten et al. 1997); GACUAG (3p arms of miR-279a to e) and AAUACU (3p arms of mir-8 and novel microRNA candidates). In addition, zygotically down-regulated genes are also enriched in target sites of mir-283-3p, mir-252-3p and mir-277-3p. Sylamer analysis (van Dongen et al. 2008) showed that of all possible 6-mer nucleotide words, motifs complementary to the seed regions of microRNAs from the above families are among the most highly and significantly enriched motifs in the 3’ UTR of the zygotically down-regulated genes (Supplemental Table 4).

**Figure 6.**
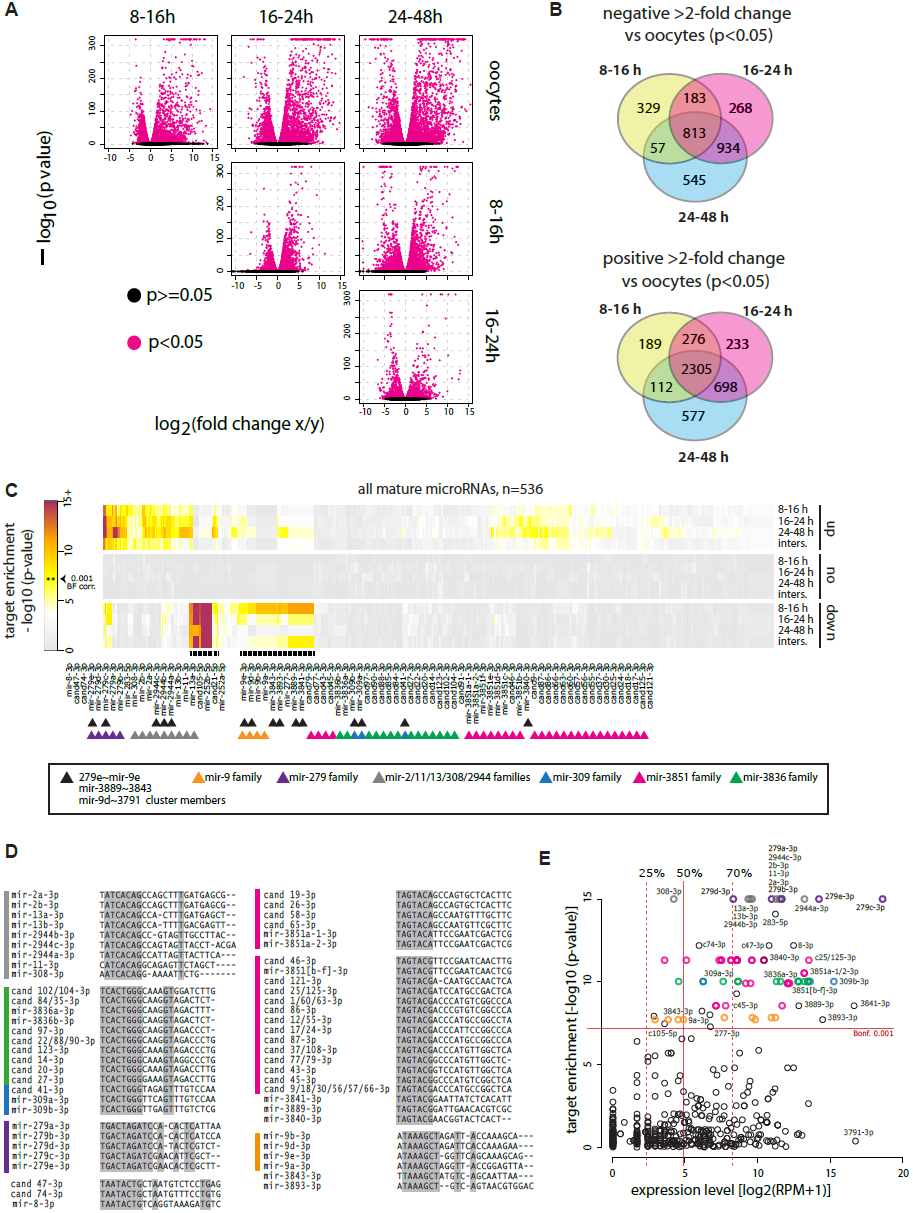
MicroRNA targeting of maternally deposited transcripts in *T. castaneum* oocytes and developing embryos. A. Volcano plots show fold change in expression (x-axis) versus differential expression p-value (y-axis) of gene expression (FPKM) in samples from *T. castaneum* oocytes, 8-16 h, 16-24h and 24-48h embryos, estimated by Cuffdiff. Transcripts with a significant fold difference between samples (p<0.05) are coloured. B. Venn diagrams showing the number and overlap between >2 fold up- and down-regulated transcripts (p<0.05) between *T. castaneum* oocytes and the three embryonic time intervals (8-16h, 16-24h and 24-48h). C. Hypergeometric p-value for mature microRNA targets enrichment predicted based on canonical site interactions among genes down-regulated (‘down’), up-regulated (‘up’) and not regulated (‘none’) from oocytes to 8-16h, 16-24h and 24-48h embryos. ‘Inters’ denotes the intersection of the three individual sets. The values underlying the heatmap representation, and full data are available as Supplemental Table 3. Labels show microRNAs with enrichment p-value < 0.001 (Bonferroni corrected). Arrowheads indicate microRNA family and clustering. D. Alignments of mature sequences of major seed families with highly enriched targets in the zygotically down-regulated gene set. Side bars indicate microRNA families colour-coded as in C. Shading indicates 100% base identity at a given position. MicroRNA with identical sequences are collapsed on a single line. E. Relationship of microRNA expression and target enrichment in the down-regulated gene set in the 8-16 h embryo. Vertical lines show the median, upper and lower quantile values, and horizontal line shows the 0.001 p-value after Bonferroni’s correction threshold. Points corresponding to the microRNAs outlined in C and D are colour-coded accordingly. Selected microRNAs are labelled.

Assessment of microRNA expression levels in the context of their targeting properties showed that the vast majority of microRNAs targeting down-regulated genes in the blastoderm are amongst the most highly expressed microRNAs at that stage (Figure 6E). These microRNAs include seed families of members of the *mir-279e∼2944c*, *mir-3889∼3843* and *mir-9d∼3791* clusters (including other mir-2/11/13, mir-279 and mir-9 loci), mir-3851, mir-3836, mir-8 and mir-283. The zygotic expression of a subset of microRNAs and the reciprocal down-regulation of their targets in the blastoderm strongly suggested that these microRNAs specifically function in maternal transcript clearance in *T. castaneum.* We also note that a different set of highly expressed and conserved microRNAs, including miR-9-5p, mir-263 and mir-276, target a substantial fraction of genes up-regulated during embryogenesis, reflecting a likely role for these microRNAs in other developmental processes (Supplemental Table 3).

Previous findings in *D. melanogaster* showed that highly expressed members of the *mir-309∼6* cluster in the early embryo, including mir-9/4/79, mir-5/6/2944, mir-3/309 and mir-279/286, are involved in maternal transcript turnover (Bushati et al. 2008). Our results suggest that these microRNA families have a conserved role in maternally deposited transcript regulation in the early embryo of holometabolous insects. Furthermore, several *Tribolium-specific* families are involved in the process, including mir-3889, mir-3840, mir-3841, and mir-3851 families. We therefore suggest that the microRNA repertoire involved is maternal transcript clearance has diverged.

## Discussion

### Quantity and modifications of maternally deposited microRNAs

Oocyte maturation is accompanied by the deposition of a large number of protein and RNA factors that not only provide trophic support, but also play important roles in the initial steps of embryonic development before the zygotic genome is activated. For instance, maternally provided morphogens are essential for the specification of body segments in insects. In *Drosophila*, the maternally deposited RNA complement also contains small RNAs, including microRNAs and piRNAs. Abundant piRNAs in *Drosophila* are thought to provide the basis of epigenetic inheritance of transposon defence (Brennecke et al. 2008). The role of maternally provided microRNAs, on the other hand, is not well understood: microRNAs can be detected in *Drosophila* oocytes, but it is unclear whether these are products generated at earlier stages of gonadal development, or if their deposition is required for subsequent events in embryonic development. For instance, genetic knockout of one of the maternally provided fruit fly microRNA clusters *mir-310∼mir-313* (Figure 4.4.4F) does not result in developmental defects in the embryo (Pancratov et al. 2013; Tsurudome et al. 2010). MicroRNAs are not present at high levels in small RNA libraries of zebrafish, *Xenopus* and mouse oocytes, and the role of maternally deposited microRNAs in these species are not well understood (Ohnishi et al. 2010; Chen et al. 2005; Watanabe et al. 2005, Lee et al. 2014a).

As in vertebrates, the cloning frequency of microRNAs in oocytes of *Tribolium* is very low compared to subsequent developmental stages. Given that the long germband developmental mode characteristic of *Drosophila* is an evolutionary derived state, it can be hypothesized that high levels of maternally deposited microRNAs is also an evolutionary innovation, uncommon in short germ-band organisms and vertebrates. However, oocytes are rich in piRNAs whose high levels of expression may be masking microRNAs in sequencing experiments. Indeed, absolute quantification of *T. castaneum* maternally deposited microRNAs shows a concentration of approximately 0.2 fmol per cell. Thus, blastoderm cells could receive thousands of copies of maternally provided microRNAs. These numbers are commensurate with previous estimates of microRNA copy numbers in mammalian cells (Bissels et al. 2009), and likely reflect physiologically-relevant levels. We propose that similarly to *T. castaneum*, maternally deposited microRNAs in other species may be present yet masked in early embryonic libraries small RNA libraries by other abundant RNAs such as piRNAs or endo-siRNAs.

A large proportion of maternally deposited mRNA products are cleared prior to or upon the activation of the zygotic genome, and this clearance is essential for the normal progression of development (reviewed in Tadros and Lipshitz 2009). The fate of maternally provided small RNAs – microRNAs and piRNAs – on the other hand is not well understood. It was recently demonstrated that the poly-A polymerase Wispy installs non-templated adenosines at the 3’-ends of specific microRNAs in oocytes in *D. melanogaster*, and this modification is likely to contribute to maternal microRNA clearance (Lee et al. 2014a). Furthermore, maternal microRNA modification is conserved in deuterostomes (Lee et al. 2014a). We have detected the highest levels of 3’-end oligoadenylated microRNAs in *Tribolium* and *Drosophila* oocytes and *Tribolium* transcriptionally inactive embryos, in agreement with the hypothesis that this modification is involved with the regulation of maternally provided microRNAs. Interestingly, only specific mature microRNAs are modified at high levels. However, Wispy-mediated adenylation is not microRNA sequence-specific, and a conserved motif between different highly adenylated microRNAs has not be identified (Lee et al. 2014). Thus, the question of how microRNA adenylation specificity is regulated *in vivo*, and its role in development, remains unsolved. Our data demonstrates that the specificity of microRNA adenylation in oocytes is similar for abundant microRNA orthologs between *Drosophila* and *Triboliium*, suggesting that this regulation is a conserved phenomenon among holometabolous insects, represented across at least ∼300 my of evolution.

### MicroRNA-mediated maternal transcript clearance in *T. castaneum*

One of the earliest events during animal development is the degradation of maternally deposited transcripts in the egg, and the activation of the zygotic genome – a process termed the maternal-to-zygotic transition (MZT) (reviewed in Tadros and Lipshitz 2009). The MZT begins with the action of pioneer transcription factors, which provide basis for the activation of the zygotic transcriptome (Iwafuchi-Doi and Zaret 2014). In both invertebrates and vertebrates, some of the earliest upregulated transcripts are microRNAs, which contribute to the degradation of maternally deposited mRNAs. However, evidence suggests that different microRNAs became involved in the MZT separately in the two clades (Bushati et al., 2008). In zebrafish, the transcription factors Nanog, Pou5f1 and SoxB1 activate the earliest zygotically expressed transcripts, including a large cluster of mir-430 paralogs (Lee et al. 2013). mir-430 sequences, in turn, promote the deadenylation and clearance of a large set of maternally deposited mRNAs (Giraldez et al. 2006). In *Drosophila*, the earliest zygotic transcriptional activation is mediated by the factor Zelda (Liang et al. 2008). Zelda up-regulates the *mir-309∼6* cluster, which is suggested to be involved in maternal transcript clearance: genetic knockout of the whole cluster results in delay of the degradation of a number of maternally provided mRNAs, which are enriched in target sites of cluster members (Bushati et al. 2008). Nonetheless, *mir-309∼6* knockout does not result in significant embryonic defects (Bushati et al. 2008). We speculate that members of the *mir-309∼6* cluster are not the only microRNAs involved in maternal transcript regulation in *Drosophila*, as other microRNAs from the same seed families – including miR-2/11/13-3p, miR-279/996-3p and mir-9 paralogs – are present in the early embryo. In line with this hypothesis, these microRNAs were recently shown to also be up-regulated early in development by Zelda (Fu et al. 2014).

We have identified a number of conserved and species-specific microRNAs in *T. castaneum* that are highly expressed in the early embryo, and target a significant fraction of down-regulated maternal transcripts during the MZT. Thus, the data suggest that, as in *Drosophila* and *D. rerio*, microRNAs are involved in the degradation of maternally deposited transcripts in this lineage. We note that the temporal antagonism of microRNAs and their targets in the early embryo is not direct evidence of microRNA-mediated degradation. Nonetheless, the observed relationships, together with the previous findings that homologous microRNAs are involved in the MZT in *Drosophila* discussed below, are strongly suggestive.

The conserved microRNAs involved in maternal transcript down-regulation in the blastoderm of the flour beetle include homologs of the *Drosophila mir-309∼6* cluster members and other microRNAs with identical seeds. Taken together, the data suggest that microRNA-mediated maternal transcript degradation by the 3p mature arms of microRNAs from the seed families ATCACA (mir-2944/5/6, mir-2/13 and mir-11), TAAAGC (mir-9/79/4 family), and CACTGG (mir-309/3 family) is a conserved feature in holometabolous insects. mir-279 family members are very highly expressed in the early embryo and show one of the strongest target enrichment values among the down-regulated transcripts in the early embryo. In addition, we observed high expression levels and target site enrichment in the zygotically down-regulated genes targeted by mir-8 and mir-283, and to a lesser extent, mir-277 and mir-252. The roles of these microRNAs in maternal transcript clearance in *Drosophila* have not been previously addressed; further studies are required to determine whether the involvement of these deeply conserved microRNAs in the MZT is conserved in insects or represents a *Tribolium-specific* co-option.

Several *T. castaneum-specific* microRNAs also target a significant fraction of the maternally deposited transcripts that decrease at the blastoderm stage. These include the multicopy microRNA families mir-3836 and mir-3851, and four hairpins encoded in the *mir-3889∼3843* and *mir-9d∼3791* clusters: mir-3889, mir-3840 and mir-3841, and mir-3893. Notably, 3p products corresponding to the seed families of mir-3851a (AGTACA) and mir-3851b to f, mir-3889, mir-3840 and mir-3841 (AGTACG) are not found in *Drosophila*, suggesting that these microRNA-target interactions are a diverged feature between these insect lineages.

Taken together with previous studies in *Drosophila*, findings in *T. castaneum* suggest that microRNA-mediated maternal transcript degradation is a conserved mechanism in holometabolous insects, but the precise microRNAs participating in this process differ somewhat between species. MicroRNAs involved in maternal transcript clearance in vertebrates (Giraldez et al. 2006; Lund et al. 2009) have no sequence similarity or common seed motifs with any of the insect microRNAs, illustrating the likelihood of convergence in this process (see below). Nevertheless, comparisons of the microRNAs in the MZT in different organisms reveal the common phenomenon of large, fast-evolving microRNA polycistrons involved in this process.

### Dynamic evolution of early expressed microRNAs

Our analysis of the genomic positions, sequences and targeting properties of the early expressed *T. castaneum-*specific microRNAs reveal complex relationships. The mir-3851 and mir-3836 families, and novel mir-8 seed family members, are found in large and diverse clusters located in multiple genomic positions. These microRNAs are not co-localized with any conserved microRNAs, but other microRNAs from these seed families, such as mir-309, have deeper evolutionary origins and are clustered with other conserved microRNAs. MicroRNA hairpins are short, and thus any putative fast-evolving sequences can diverge to the point at which they cannot be confidently identified as homologs. On the other hand, the formation of microRNAs with identical seeds (microRNA convergence) may be common, as the microRNA seed region is very short, and new hairpins often emerge *de novo* in animal genomes. Despite their high degree of divergence in terms of encoded hairpin copy number, family and sequence, the *T. castaneum-specific* microRNA clusters display a very similar temporal expression pattern spanning only a few rounds of cell division after the initiation zygotic transcription. We propose that newly-emerged microRNAs with convergent seeds and similar expression patterns to existing microRNAs are more likely to be retained, as they ‘mimic’ the existing microRNA and thus do not cause significant transcriptome perturbations by down-regulating new transcripts. In the light of this hypothesis, one explanation for the origin of the mir-3836 and mir-3851 clusters is that their founding hairpins emerged in early activated regions from random hairpins with similar seeds to existing microRNAs, and subsequently duplicated and diversified. Alternatively, if microRNAs with identical seeds are considered to be highly diverged paralogs, we can speculate that the mir-3836 and mir-3851-encoding clusters, and the three clusters encoding members of the conserved mir-5/6/2944, mir-9/4, mir-279/286 and mir-309/3 families, have common origins but have significantly diverged via multiple duplications, rearrangements and losses (including the acquisition of a mir-8 paralog that rapidly diverged). These scenarios of cluster evolution are not mutually exclusive: it is likely that some seed families are evolutionarily related, while others emerged by convergence. Either way, the evolutionary patterns of the early expressed microRNAs are uniquely dynamic.

Our previous work demonstrated that one characteristic of the early *Drosophila* embryo is high levels of fast-evolving microRNAs (Ninova et al. 2014b). Data from *T. castaneum* now suggests that the early embryonic expression of fast-evolving and evolutionarily younger microRNAs is not restricted to *Drosophila*, but represents a conserved feature of holometabolous insects. Studies in other organisms have also suggested that early embryogenesis is permissive or robust to evolutionary change in the transcriptome compared to later stages of development: in *Drosophila*, vertebrates, and plants the early embryonic transcriptome is on average younger, faster evolving, and characterized with higher variation in orthologous gene expression (Kalinka et al. 2010; Heyn et al. 2014; Quint et al. 2012; Domazet-Lošo and Tautz 2010). Even though the underlying causes of this phenomenon are elusive, our results suggest that the apparent flexibility of the molecular networks active in early development also impacts the evolution of the microRNA complement expressed at that stage.

## Methods

### Animal husbandry and sample collection

*T. castaneum* wild-type adults (Michael Akam, University of Cambridge) were reared and embryos were collected following a standard protocol (The Beetle Book (http://wwwuser.gwdg.de/~gbucher1/tribolium-castaneum-beetle-book1.pdf) at 25°C. Unfertilized eggs were obtained from virgin females isolated at the pupal stage that were allowed to lay for 4 hours. *D. melanogaster* (w^1118^) and wild type *D. virilis* unfertilized eggs were collected from virgin females allowed to lay for 4 hours on apple juice agar plates supplemented with yeast paste, at 25°C. Samples were dechorionated in 0.5-1% hypoclorite solution (Sigma) for 1-2 min, and thoroughly washed.

### Embryo fixation, immunohistochemistry and *in situ* hybridization

Dechorionated embryos were fixed and devitellinated using a standard protocol. Whole-mount fluorescent *in situ* hybridization with DIG-labelled antisense RNA probes and antibody staining procedures were performed according to the protocol in (Kosman et al. 2004), but omitting the proteinase K treatment step. RNA synthesis templates were amplified from *T. castaneum* genomic DNA using the following primer pairs: CGAAATTTCTGCCAGGTCCA/ AAGAACGCGCATTTTACAAA for the candidate-20 region; TGCTTCATAATACCAGACACTCC/ GACCGGGCAGAAATTTCGAA for the candidate-25 region; GAACCAATCAGAAAAGGGGTACT/ CAAGCAGGCTAGGAGACAGA for the candidate-75∼candidate-40 region; TCCCAACGCTGTATAAGGCA/AAATTGCTTCTTCTGCGGCA for the candidate-85 region; CCTGCAAATGAAGAGTGGGG/TAGGTCGCCCAGATGAACAG for the *mir-9d-3791* cluster region and CAGTGCTGGATTTGGAAACA/TCCACTAAAGCCACATCAACA for *mir-3889∼3843* cluster region. We used sheep anti-DIG (Roche), mouse anti-RNA Polymerase II H5 (Covance) and mouse anti-engrailed (4D9, DSHB) primary antibodies, anti-sheep Alexa Fluor®555 and anti-mouse Alexa Fluor®488 secondary antibodies (Life Technologies), and DAPI mounting media (ProLong Gold, Invitrogen). Images were visualized by confocal microscopy on an Olympus FV1000, and image stacks were processed with Fiji (Schindelin et al. 2012).

### RNA isolation and deep sequencing libraries

For *T. castaneum* small RNA developmental profiles, we collected embryonic samples representing 0-5 h (in two replicates), 8-16 h, 16-20 h, 20-24h, 24-34 h, 34-48 h and 2-6 d time intervals at 25°C. Small RNA libraries from unfertilized eggs from *T. castaneum*, *D. virilis* and *D. melanogaster* were prepared in two biological replicates. For whole transcriptome RNA sequencing, we used two biological replicates of *T. castaneum* unfertilized eggs, 8-16 h, 16-24 h and 24-48 h embryos.

An aliquot from each sample was preserved for microscopic visualization. The remaining embryos were disrupted in Trizol (Life Technologies) and total RNA was extracted according to the manufacturer’s instructions, and adding 0.5 μl Glycoblue (Life Technologies) as a co-precipitant. For total RNA from unfertilized eggs, we manually picked 30 eggs per sample immediately after collection, and RNA pellets were re-suspended in 30 μl cocktail containing 0.1 nM cel-miR-35-3p, 0.01 nM cel-miR-56-3p and 0.001 nM cel-miR-230-3p 5’ phosphorylated synthetic oligonucleotides as spiked-in normalization controls. Small RNA libraries were constructed from total RNA using the Illumina TruSeq® Small RNA Sample Prep Kit, and whole transcriptome libraries were prepared using the Illumina TruSeq® Stranded mRNA Sample Prep Kit according to the manufacturer’s instructions. Libraries were assessed using the Agilent 2200 TapeStation, and sequencing was performed on the Illumina MiSeq (embryonic small RNA sequencing), or the Illumina HiSeq 2000 (oocytes small RNA sequencing and whole-transcriptome sequencing) platforms in the University of Manchester Genomic Technologies facility, generating respectively 50 bp and paired-end 101 bp reads. Data was submitted to the GEO database under accession number GSE63770.

### Small RNA sequencing data analysis and microRNA prediction

Adapter sequences were trimmed from the *T. castaneum* small RNA reads using the Cutadapt tool (http://code.google.com/p7cutadapt/), retaining reads longer than 16 nucleotides. The trimmed reads were first filtered against *T. castaneum* tRNA genes predicted using tRNAscan-SE (v1.3, Lowe and Eddy 1997), and then mapped to the latest version of the *T. castaneum* genome assembly (r4.0) using Bowtie (v1.0, Langmead et al. 2009) with the following parameters: -v 1 −a --best −strata −m 5. Mapped reads were used as input to two independent microRNA discovery methods – mirdeep2 (Friedländer et al. 2008) and an implementation of the method described in (Marco et al. 2010). Newly discovered microRNAs were submitted to miRBase (Kozomara and Griffiths-Jones 2014). Read counts for known and newly annotated mature *T. castaneum* microRNA sequences were computed using a custom perl script, correcting for mapping to multiple locations. Genomic origins of non-microRNA reads (intragenic versus intergenic) were inferred by intersection with the *T. castaneum* protein-coding gene annotations accompanying the 4.0 genome release. Expression was normalized as reads per million mapped to the genome.

*D. melanogaster* and *D. virilis* oocyte small RNA libraries were trimmed in a similar manner, and reads mapping to rRNAs, tRNAs, snRNAs and snoRNAs were filtered out. Reads were then mapped to the *D. melanogaster* r5.4 and *D. virilis* r1.2 genomes, respectively, and annotated based on miRBase (v20, Kozomara and Griffiths-Jones 2014) and the corresponding protein-coding gene annotations for each genome release.

For 3’ non-template end additions, reads that did not map to the genome with 0 mismatches were sequentially trimmed by 1 nucleotide from their 3’-end and remapped to the genome using Bowtie with the following parameters: -a --best --strata -v 0. Upon each iteration, perfectly mapping reads were retained. Resulting trimmed sequences that correspond to full-length microRNAs were then inspected for the type of 3’-end addition by comparison with the original read sequence using a custom perl script.

### MicroRNA evolutionary conservation

*T. castaneum* microRNAs were grouped into families based on best blastn hits (- word_size=4, Altschul et al. 1990) and manual inspection and editing of the resulting alignments using RALEE (Griffiths-Jones 2005). Curated alignments were used to build covariance models, and these models were searched against the genomes of *Dendroctonus ponderosae*, *D. melanogaster*, *D. virilis*, *A. mellifera* and *B. mori* using INFERNAL (Nawrocki et al. 2009) with an E-value cutoff of 1. Hits scoring below this threshold were added to previous alignments and inspected for hairpin folding and homology.

### RNA sequencing data analysis and differential expression

Paired-end transcriptome data was mapped to the *T. castaneum* genome (r4.0) using Tophat (Trapnell et al. 2009) with default parameters and supplying the currently available protein-coding gene annotations (iBeetle, http://bioinf.uni-greifswald.de/tcas/genes/annotation/). Gene expression counts were obtained using htseq-count (Anders et al. 2014) and differential gene expression between oocytes, 8-16 h, 16-24 h and 24-48 h embryos was assessed using the DESeq R package (Anders and Huber 2010). Gene expression changes with p-values smaller than 0.05 after Benjamini-Hochberg correction were considered as significant. We further validated these results using the Cuffdiff program from the Cufflinks package (Trapnell et al. 2010) as an alternative approach to estimate gene differential expression (Supplemental Table 3). Downstream analyses were performed using custom R-scripts.

### MicroRNA target predictions and enrichment analyses

MicroRNA target sites in the annotated 3’ UTR of *T. castaneum* protein-coding genes (r4.0, iBeetle Database, http.//ibeetle-base.uni-goettingen.de/’) were predicted using two independent target prediction algorithms with the default parameters – an implementation of the canonical site target pairing as described in (Bartel 2009), provided by Antonio Marco, and miRanda (Enright et al. 2003). In the analyses presented in the main text, we considered differential expression on the gene level, and thus when multiple transcripts per gene are present (7% of all annotations), overlapping UTRs were merged, and non-overlapping UTRs were concatenated thereby pooling microRNA target sites of different transcripts together; we note that considering individual transcripts separately produces very similar results (data not shown). Target enrichment was calculated based on the hypergeometric distribution (phyper function in R and sylamer (van Dongen et al. 2008), and multiple testing was accounted for using the Bonferroni correction for 536 microRNAs tested for target enrichment in 12 gene sets (up-, down- and not-regulated genes between oocytes and 8-16 h, 16-24h, 24-48h embryos and all stages).

## Data Access

All RNA sequencing data are deposited in the Gene Expression Omnibus (GEO) database under accession number GSE63770.

## Acknowledgements

We thank Mario Stanke and Gregor Bucher for providing the *T. castaneum* r4.0 release and protein-coding gene annotations.

## Author contributions

MN, MR and SGJ conceived the study, designed the experiments and wrote the manuscript. MN performed all experiments and data analyses, and drafted the manuscript.

## Supplemental Material

Supplemental_Figures.pdf

Supplemental_Text.doc (including Supplemental Figure Legends)

Supplemental_Table1.xlsx

Supplemental_Table2.xlsx

Supplemental_Table3.xlsx

Supplemental_Table4.xlsx

